# Pregnancy alters innate immune responses to Zika virus infection in the genital tract

**DOI:** 10.1101/828731

**Authors:** Kelsey E. Lesteberg, Dana S. Fader, J. David Beckham

## Abstract

Recent outbreaks of Zika virus (ZIKV) have been associated with birth defects, including microcephaly and neurological impairment. However, the mechanisms which confer increased susceptibility to ZIKV during pregnancy remain unclear. We hypothesized that poor outcomes from ZIKV infection during pregnancy are due in part to pregnancy-induced alteration of innate immune cell frequencies and cytokine expression. To examine the impact of pregnancy on innate immune responses, we inoculated pregnant and non-pregnant female C57BL/6 mice with 5×10^5^ FFU of ZIKV intravaginally. Innate immune cell frequencies and cytokine expression were measured by flow cytometry at day 3 post infection. Compared to non-pregnant mice, pregnant mice exhibited higher frequencies of uterine macrophages (CD68+) and tolerogenic dendritic cells (CD11c+ CD103+ and CD11c+ CD11b+). Additionally, ZIKV-infected pregnant mice had lower frequencies of CD45+ IL-12+ and CD11b+ IL-12+ cells in the uterus and spleen. These data show that pregnancy results in an altered innate immune response to ZIKV infection in the genital tract of mice and that pregnancy-associated immune modulation may play an important role in the severity of acute ZIKV infection.

**Importance:** Pregnant females longer duration that viremia following infection with Zika virus but the mechanism of this is not established. Innate immune cellular responses are important for controlling virus infection and are important for development and maintenance of pregnancy. Thus, the acute immune response to Zika virus during pregnancy may be altered so that the pregnancy can be maintained. To examine this interaction, we utilized a mouse model of Zika virus infection during pregnancy using intravaginal inoculation. We found that following Zika virus infection, pregnant mice exhibited increased expression of tolerant or non-inflammatory dendritic cells. Additionally, we found that pregnant mice have significantly depressed ability to secrete the cytokine IL-12 from innate immune cells in the uterus and the spleen while maintaining MHCII expression. These findings show that pregnancy-induced changes in the innate immune cells are biased towards tolerance and can result in decreased antigen-dependent stimulation of immune responses.

## Introduction

Zika virus (ZIKV) is a neurotropic flavivirus originally isolated from a febrile rhesus macaque in the Zika Forest of Uganda (1). Although ZIKV was first identified in humans in 1952, only sporadic infections occurred in humans until 2007, when the first major outbreak was reported on Yap Island (2). Since its emergence, infections have been reported in Africa, Asia, the Pacific Islands, and the Americas. ZIKV is transmitted to humans predominantly through bites from *Aedes* mosquitos, but infections after sexual contact and blood transfusions have also been reported (3, 4). The majority of infected individuals are asymptomatic, with some infections causing mild symptoms such as fever, rash, conjunctivitis, muscle and joint pain, malaise, and headache (5). However, ZIKV infections in French Polynesia in 2013 and 2014 were linked to an increase in Guillain-Barré syndrome in adults (6). In 2015, a ZIKV outbreak in Brazil was associated with a marked increase in microcephaly in infants born to acutely infected mothers (7-9). Other reports also show that ZIKV infection in fetuses may cause a spectrum of disease from severe microcephaly to more subtle brain and developmental abnormalities together referred to as “congenital ZIKV syndrome” (8, 10-12). Multiple studies have provided additional evidence of vertical transmission of ZIKV from infected pregnant mothers to the fetus (9, 13). Vertical transmission of ZIKV often occurs following periods of prolonged maternal viremia, and this is supported by data from both human studies and nonhuman primate models of congenital ZIKV infection (14, 15).

The mechanisms underlying the increased severity of ZIKV infection during pregnancy remain understudied. During pregnancy, women are at increased risk for infection and increased severity of infection with ZIKV and several other pathogens, including listeria, cytomegalovirus (CMV), herpes simplex virus (HSV), influenza virus, and HIV (16-21). In general, successful pregnancy relies on tolerance of the maternal immune system towards the semi-allogeneic fetus, which is often referred to as immunotolerance. This results in changes at multiple levels of the maternal immune system. For example, human natural killer (NK) cells lose their cytotoxic abilities and instead take on a supportive role during pregnancy (22). Additionally, the decidua contains an abundance of regulatory T cells (Tregs) during early pregnancy, which maintain tolerance, prevent inflammation, and promote implantation of the embryo (23-25). Many of these changes are linked to the induction of pregnancy hormones. Several studies have suggested that human chorionic gonadotropin (HCG) plays a role in the recruitment of Tregs to the maternal-fetal interface and promotes the generation of tolerogenic dendritic cells (DCs) (26, 27). These pregnancy-induced immune changes impact susceptibility to several pathogens. For example, changes in the immune response during pregnancy result in progesterone-dependent increased susceptibility to HSV2 in the genital tract of mice, resulting in lower HSV2-specific IgG and IgA responses in the genital tract following infection (28). Additionally, influenza infection in pregnant ferrets results in decreased total CD8+ T-cells and decreased H1N1-specific B-cell responses compared to non-pregnant ferrets (29).

While different stages of pregnancy clearly modulate adaptive immune responses, less is known about pregnancy-induced innate immune responses during viral infection. In a pregnant mouse model of influenza infection, Cox-2, PGE2, and PGF2α were increased, resulting in remodeling of the placental architecture, preterm labor, impaired fetal growth, and increased fetal and maternal mortality and morbidity (18). In another study of late stage pregnancy, viral infection of the placenta triggered an inflammatory response, including the secretion of IL-1, IL-6, IL-8, and TNFα, and fetal abnormalities in the absence of direct fetal infection (30, 31). A study of ZIKV-infected mothers reported the presence of interferon gamma-inducible protein-10 (IP-10), IL-6, IL-8, monocyte chemoattractant protein-1 (MCP-1), vascular endothelial growth factor (VEGF), and granulocyte-colony stimulating factor (G-CSF) in the amniotic fluid of mothers whose infants were born with microcephaly (32). Another study found that IP-10, CCL5, IL-9, interferon gamma (IFNγ), IL-7, IL-5, and IL-1ra were upregulated in the plasma of acutely infected individuals compared to healthy donors (33). In the recovery phase, IL-12p70 and basic fibroblast growth factor (FGF) were found to be upregulated (33). However, it is unclear how the immune response to ZIKV is impacted by pregnancy, especially in early pregnancy.

Few studies have examined the effect of pregnancy on innate immune responses in the genital tract at the early stages of pregnancy prior to placental formation. Previous studies have shown that ZIKV infections in the first trimester of pregnancy confer a greater risk of microcephaly compared to second and third trimester infections (34). Since the severity of ZIKV congenital disease increases with infection during early stages of pregnancy, we examined pregnancy-associated changes in the innate immune response during early pregnancy. In order to evaluate the innate immune cellular response in the genital tract, we utilized an immune competent murine model of ZIKV intravaginal inoculation which has been described previously (35). Following intravaginal inoculation of ZIKV at embryonic day 4.5 (E4.5), we found that pregnant mice exhibited increased frequencies of tolerogenic DCs (CD11c+ CD103+, CD11c+ CD11b+) in the uterus and a higher frequency of uterine macrophages (CD68+) compared to ZIKV-inoculated non-pregnant mice. Additionally, ZIKV-infected pregnant mice exhibited lower frequencies of CD45+ IL-12+ cells and CD11b+ IL-12+ in the uterus and spleen. Taken together, these results suggest that pregnancy alters the local innate immune response to ZIKV infection, which may decrease immune control of acute viral infection.

## Results

### Intravaginal ZIKV infection in C57BL/6 mice

To generate a mouse model of ZIKV infection during pregnancy, 8-week-old, female C57BL/6 mice were injected with 2.5 international units (iu) pregnant mare serum gonadotropin (PMSG) by intraperitoneal (ip) inoculation, ip injected with 2.5 iu human chorionic gonadotropin (HCG) 48 hours later, and finally mated with male mice overnight (16 hours). The males and females were separated the following morning (E0.5). At E4.5, the female mice were inoculated with 5×10^5^ FFU of ZIKV (PRVABC59) or mock inoculum. Intravaginal washes and tissue harvests for analysis of virus and immune parameters, respectively, were performed at specific time points post-infection (**Fig. 1A**). Pregnancy rates ranged from 30-80% with this approach, allowing for prospective cohort analysis of both pregnant and non-pregnant mice that were treated at the same time with the same hormones prior to ZIKV infection. At 48 hours post-infection, ZIKV PCR of vaginal wash fluid revealed evidence of ZIKV RNA in the genital tract of both pregnant and non-pregnant mice (**Fig. 1B**). The levels of ZIKV RNA did not differ between the pregnant and non-pregnant mice. ZIKV PCR from vaginal washes at days 1, 2, and 3 post-infection revealed peak values at day 2 post-infection, and ZIKV RNA was not detected in fetal tissues at day 3 or 6 post-infection (data not shown). The weights of the mice did not change significantly during infection (data not shown), and the numbers of fetuses in both the mock and ZIKV-infected pregnant mice were similar (**Fig. 1C**).

**FIG 1.**
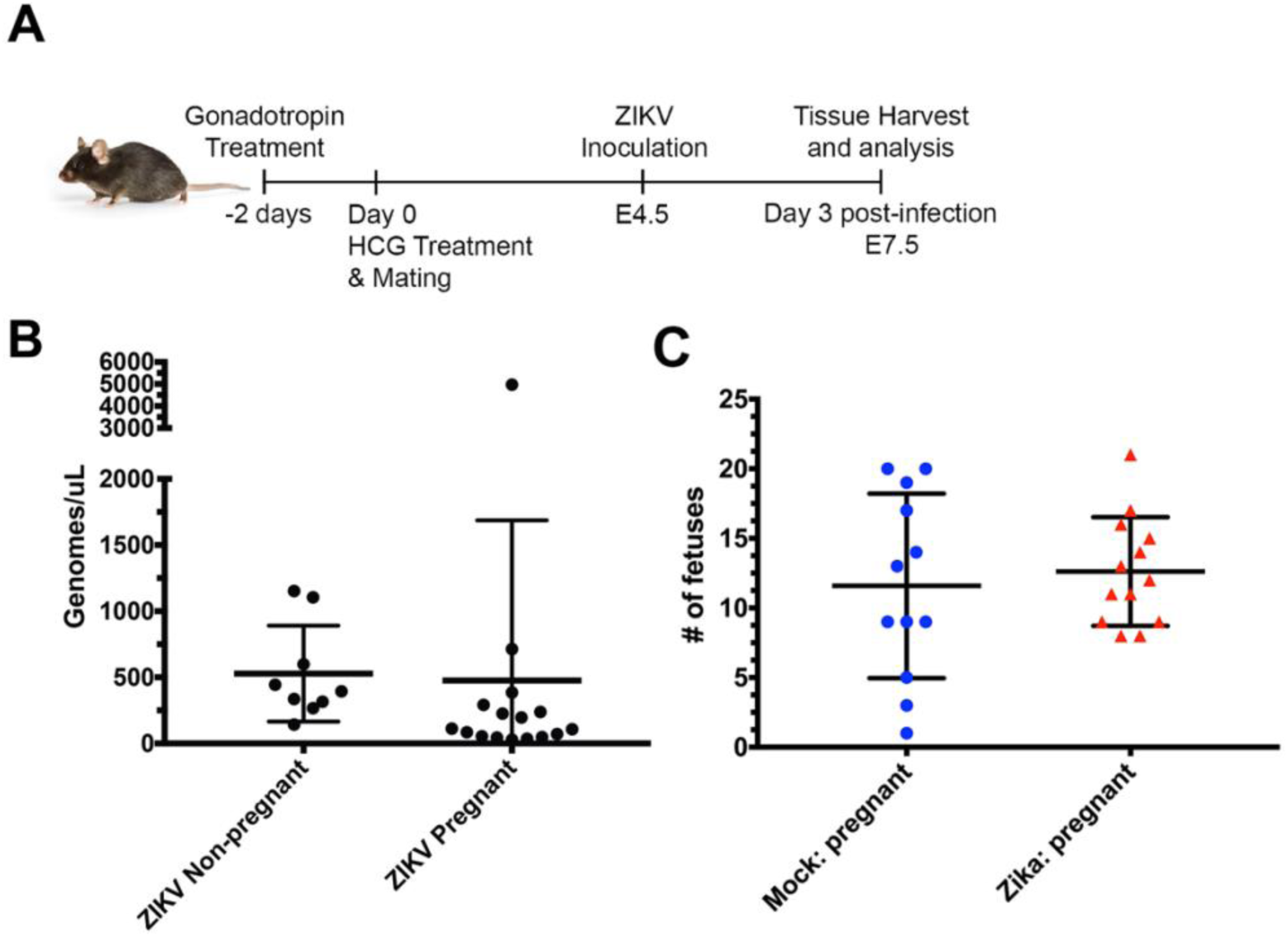
Intravaginal ZIKV infection in C57BL/6 mice. **A)** Experimental methodology: female C57BL/6 mice were treated with gonadotropins, mated, and then infected intravaginally with ZIKV at embryonic day 4.5 (E4.5). Spleen and uterine tissues were harvested at day 3 post infection (E7.5). **B)** Vaginal lavages were performed 48 hours post infection, and ZIKV RNA was detected by qPCR. N=25. **C)** Fetuses were dissected from intact uteruses from ZIKV-infected (red triangles) and mock-infected (blue circles) pregnant mice. N=25.Error bars represent the mean ± standard deviation.

### Pregnancy-induced changes in splenic innate cellular response during ZIKV infection

Nex, we analyzed the innate cellular immune responses in the pregnant and non-pregnant mice at 3 days post-infection, immediately following peak ZIKV infection of the genital tract. Following mock and ZIKV intravaginal inoculation, non-pregnant and pregnant mice were euthanized at day 3 post-infection (E7.5), and spleen and uterine tissue were analyzed by flow cytometry. We measured the frequencies of cells expressing several markers of innate immune cells, including CD68 (macrophages), CD11b (expressed on monocytes, macrophages, and DCs), CD11c (expressed on DCs, monocytes, macrophages, and granulocytes), Ly6C (expressed on macrophages, monocytes, and neutrophils), and CD103 (expressed on certain DC subsets). In the splenic tissue, ZIKV infection during pregnancy resulted in a significantly increased frequency of CD45+ CD68+ macrophages compared to ZIKV-infected non-pregnant mice (**Fig. 2A,** p= 0.0095, ANOVA=0.0107). There was no significant difference in the frequency of CD45+ CD68+ macrophages between the mock-inoculated pregnant and non-pregnant mice (p= 0.5849, ANOVA=0.5849). Mock and ZIKV-infected pregnant and non-pregnant mice exhibited similar frequencies of CD45+ CD11b+ cells (**Fig. 2B**; p= 0.984 mock non-pregnant vs mock pregnant, p= 0.9992 ZIKV non-pregnant vs ZIKV pregnant; ANOVA= 0.7906) and CD45+ CD103+ cells (**Fig. 2E**; p= 0.8707 mock non-pregnant vs mock pregnant, p= 0.9937 ZIKV non-pregnant vs ZIKV pregnant; ANOVA= 0.8585) in the spleen. However, ZIKV-infected pregnant mice exhibited increased frequencies of CD45+ CD11c cells in the spleen compared to infected non-pregnant mice (**Fig. 2C**, p= 0.0109, ANOVA=0.0104). Additionally, ZIKV infection during pregnancy resulted in a significantly decreased frequency of CD45+ Ly6C+ cell populations compared to ZIKV-infected, non-pregnant mice (**Fig. 2D**, p= 0.042, ANOVA=0.0319). In comparison, mock-inoculated pregnant and non-pregnant mice exhibited no significant changes in the frequency of splenic CD45+ Ly6C+ cells (p= 0.4801). In mice, Ly6C expression in CD11b+ monocytes distinguishes pro-inflammatory monocytes from anti-inflammatory patrolling monocytes which participate in tissue repair, with the pro-inflammatory group having high expression of Ly6C (Ly6C hi) and the anti-inflammatory group having low expression of Ly6C (Ly6C lo). Pregnant and non-pregnant infected and non-infected mice had similar frequencies of CD11b+ Ly6C hi (ANOVA= 0.2511) and Ly6C lo cells, although there was a significant difference in Ly6C lo frequencies between the pregnant mock and non-pregnant infected mice (**Fig. 2G**, p= 0.0122, ANOVA=0.0147). These data show that Ly6C expression on CD11b+ monocytes is decreased in the spleen during pregnancy despite acute viral infection.

**FIG 2.**
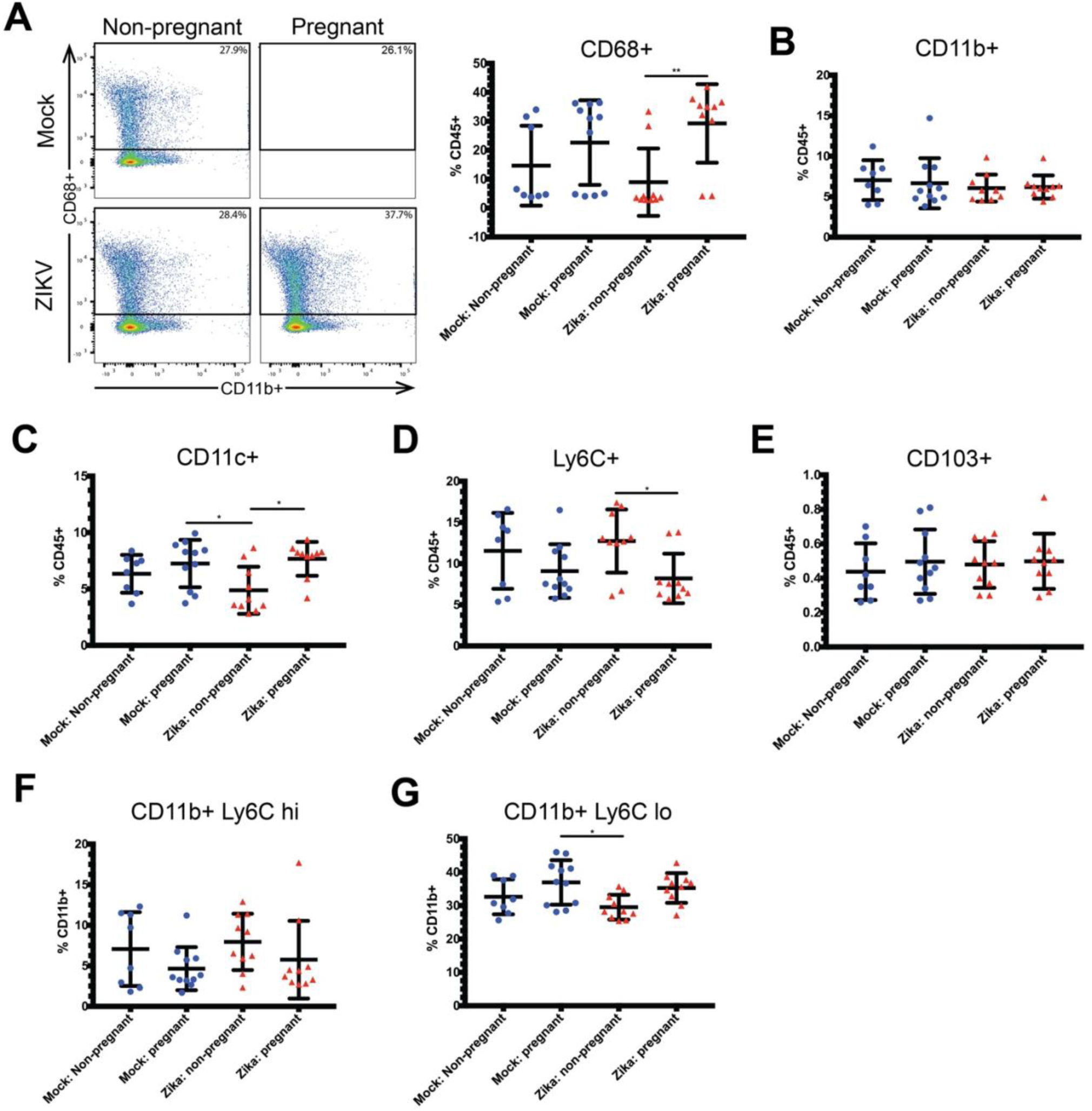
Changes to the peripheral immune response in the spleen during pregnancy and ZIKV infection. Flow cytometry was performed on the splenic immune cells of ZIKV-infected (red triangles) and mock-infected (blue circles) pregnant and non-pregnant mice, and frequencies of CD68+ (**A**), CD11b+ (**B**), CD11c+ (**C**), Ly6C+ (**D**), CD103+ (**E**), CD11b+ Ly6C high (hi) (**F**), and CD11b+ Ly6C low (lo) (**G**) cells were measured. Representative pseudocolor plots showing gating for CD11b are shown in (A). *p<0.05, **p <0.01; one-way ANOVA and Tukey’s multiple comparisons test. N=39. Error bars represent the mean ± standard deviation.

### Pregnant mice have higher frequencies of CD68+ macrophages in uterine tissue

Next, we evaluated innate immune cell frequencies in the uterus at day 3 post infection. Following infection, both ZIKV-infected and mock-infected pregnant mice exhibited increased frequencies of uterine CD45+ CD68+ macrophages compared to non-pregnant mice (**Fig. 3A**; p= 0.0055 mock non-pregnant vs. mock pregnant, p= 0.0004, ZIKV non-pregnant vs. ZIKV pregnant; ANOVA <0.0001). Despite the changes in CD68+ cell frequencies, we found no significant changes in the frequencies of CD45+ CD11b+ (**Fig. 3B**, ANOVA= 0.057), CD45+ CD11c+ (**Fig. 3C**, ANOVA= 0.7392), CD45+ Ly6C+ (**Fig. 3D**, ANOVA= 0.1022), CD11b+ Ly6C hi (**Fig. 3E**, ANOVA= 0.1915), or CD11b+ Ly6C lo (**Fig. 3F**, ANOVA= 0.0794) cell populations when comparing mock and ZIKV-infected non-pregnant and pregnant mice.

**FIG 3.**
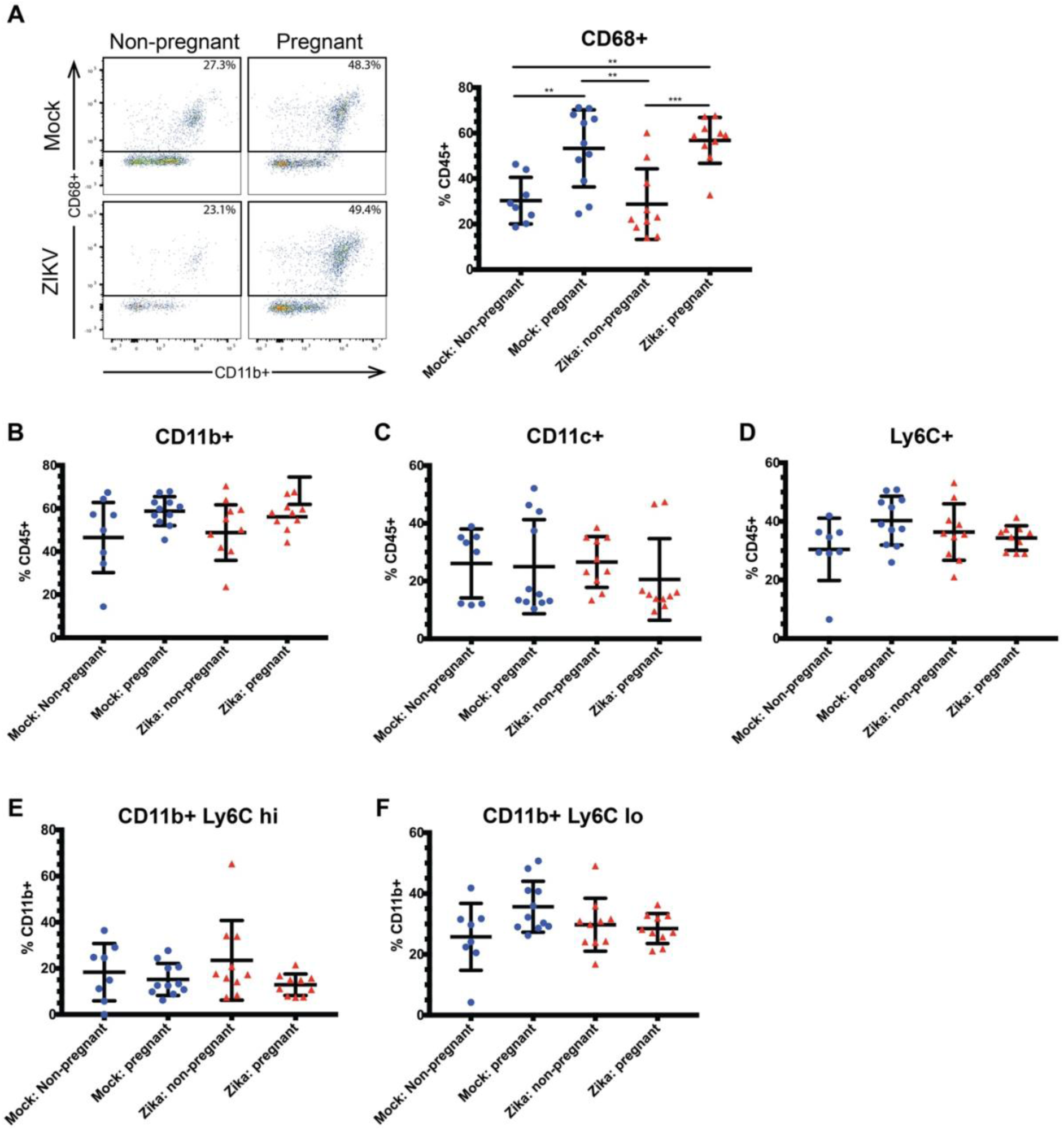
Pregnant mice have higher frequencies of uterine CD68+ macrophages. Flow cytometry was performed on the uterine immune cells of ZIKV-infected (red triangles) and mock-infected (blue circles) pregnant and non-pregnant mice. Frequencies of CD68+ (**A**), CD11b+ (**B**), CD11c+ (**C**), Ly6C+ (**D**), CD11b+ Ly6C hi (**E**), and CD11b+ Ly6C lo (**F**) cells were measured. Left panel, A) representative pseudocolor plots showing gating for CD68+ cells. **p<0.01, ***p<0.001; one-way ANOVA and Tukey’s multiple comparisons test. N=40 for (B), N=39 for all other panels. Error bars represent the mean ± standard deviation.

### Pregnant mice have higher frequencies of tolerogenic DCs

Since infiltrating macrophages didn’t express pregnancy-associated changes in activation in the uterine tissue following acute viral infection, we evaluated dendritic cells (DCs) for evidence of pregnancy-induced changes in activation during acute viral infection. DCs are important antigen presenting cells which coordinate the innate immune response and support the development of adaptive immune responses. Several lines of evidence suggest that uterine dendritic cells take on a tolerogenic phenotype during pregnancy (36, 37). Tolerogenic DCs are potent secretors of anti-inflammatory mediators such as IL-10 and weak producers of pro-inflammatory cytokines including IL-12 and TNFα (38, 39). Two types of tolerogenic DCs are present in the murine uterus: those positive for CD103 (CD11c+ CD103+) and those double-positive for CD11c and CD11b (CD11b+ CD11c+) (40). Despite what is known about these cells during pregnancy, little is known about DC activation during acute viral infection of the genital tract. Therefore, we evaluated uterine and splenic tissue for changes in tolerogenic DC populations after ZIKV infection. At 3 days post-infection, ZIKV-inoculated pregnant mice exhibited significantly increased frequencies of uterine CD45+ CD11b+ CD11c+ cells compared to non-pregnant mice (**Fig. 4A**, p=0.0004, ANOVA= 0.0001). Similarly, ZIKV-infected pregnant mice had a significantly greater frequency of splenic CD11b+ CD11c+ cells compared to mock-inoculated mice (**Fig. 4B**; p=0.195 mock non-pregnant vs. ZIKV pregnant, p=0.015 mock pregnant vs. ZIKV pregnant; ANOVA=0.0074). Within the uterine CD11b+ CD11c+ population, ZIKV-inoculated pregnant mice also exhibited a significant increase in MHCII expression (**Fig. 4A**, p=0.0081, ANOVA= 0.0039). Despite the change in MHCII expression, both the mock and ZIKV-inoculated pregnant mice exhibited markedly decreased CD11b+ CD11c+ MHCII+ CD86+ cell frequencies (**Fig. 4C**; p=0.017 mock non-pregnant vs. mock pregnant, p=0.0007 ZIKV non-pregnant vs. ZIKV pregnant; ANOVA <0.0001). CD86 expression in the splenic CD11b+ CD11c+ population did not differ between groups (ANOVA, p=0.2215). While frequencies of CD11b+ CD11c+ MHCII+ cells increased with pregnancy, we found that frequency of IL10 expression decreased during pregnancy, although not significantly during ZIKV infection (**Fig. 4D**, p= 0.0311 mock non-pregnant vs. pregnant, p=0.306 ZIKV non-pregnant vs. pregnant; ANOVA= 0.016). These data show that pregnancy induces expression of tolerogenic DCs (CD45+ CD11b+CD11c+) that express increased MHCII with ZIKV infection during pregnancy while decreasing expression of CD86. Moreover, IL-10 expression is significantly decreased during this early stage of pregnancy in mock infected animals, but, IL-10 is not significantly suppressed in pregnant mice during ZIKV infection. These data show that during pregnancy, ZIKV infection induces increased expression of tolerogenic signals (decreased CD86 and less suppression of IL-10) on CD11b+CD11c cells.

**FIG 4.**
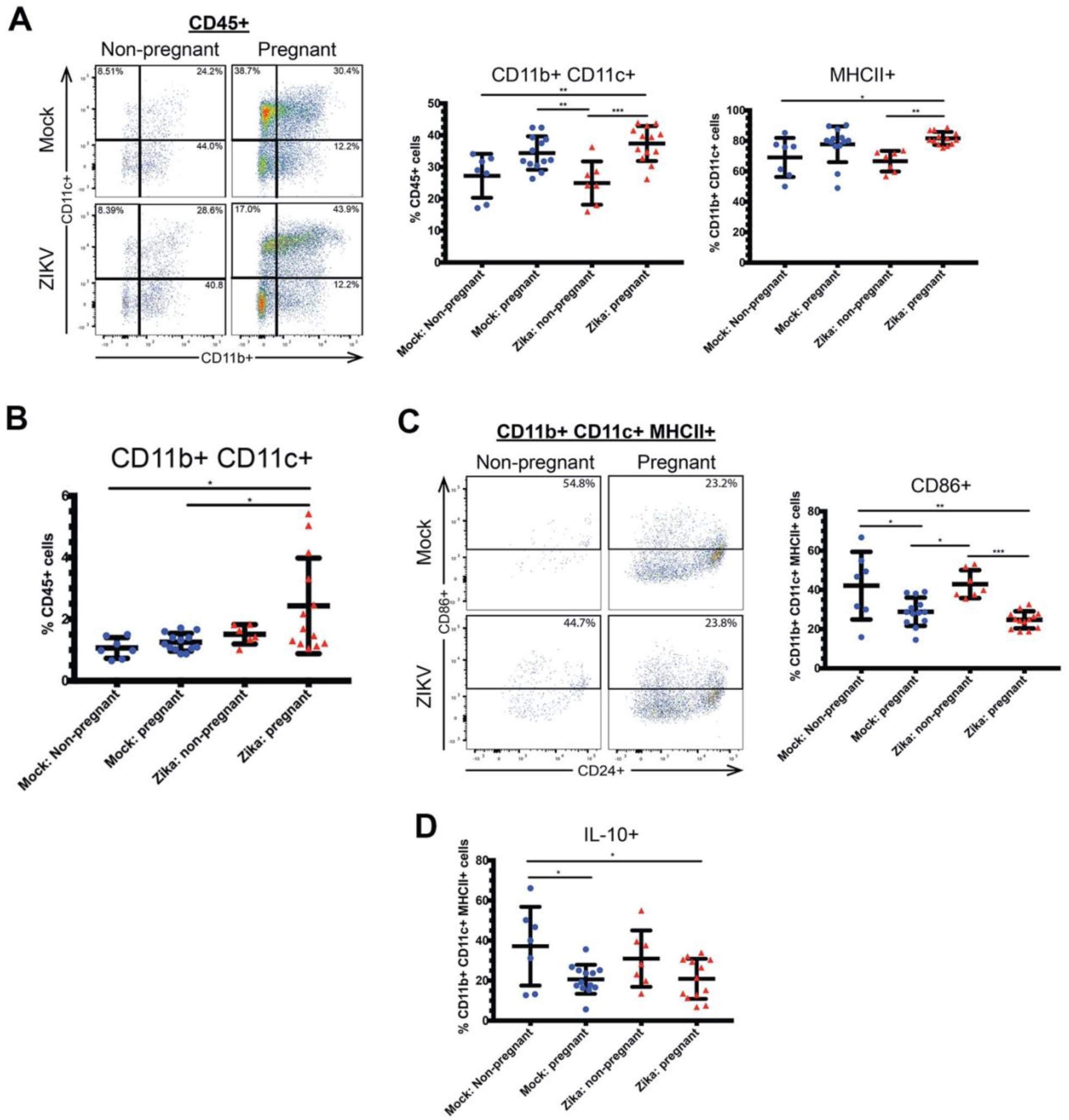
Pregnant mice have higher frequencies of CD11b+ CD11c+ tolerogenic dendritic cells in the uterus. Flow cytometry was performed on the uterine immune cells of ZIKV-infected (red triangles) and mock-infected (blue circles) pregnant and non-pregnant mice. **A)** Frequencies of CD11b+ CD11c+ cells and CD11b+ CD11c+ MHCII+ cells in the uterus and (**B**) frequencies of CD11b+ CD11c+ cells in the spleen. **C)** CD86+ CD11b+ CD11c+ MHCII+ cells, and **D)** IL-10+ CD11b+ CD11c+ MHCII+ cells were measured. Representative flow cytometry plots are also shown in A (CD11b+ CD11c+ cells within the CD45+ population) and B (CD86+ cells within the CD11b+ CD11c+ MHCII+ population). *p<0.05, **p<0.01, ***p<0.001; one-way ANOVA and Tukey’s multiple comparisons test. N=40. Error bars represent the mean ± standard deviation.

Similar to CD11b+ CD11c+ cells, we found that ZIKV-inoculated, pregnant mice exhibited a significant increase in frequency of uterine CD11c+ CD103+ cells (**Fig. 5A**, p= 0.0004 ZIKV non-pregnant vs. ZIKV pregnant, ANOVA= 0.0001). ZIKV infection in non-pregnant mice resulted in a suppression of this tolerogenic cell population but pregnant mice still exhibited high levels of CD11c+ CD103+ cells despite ZIKV infection. Within the CD11c+ CD103+ population, there was no significant change in the CD86+ (ANOVA=0.2347) or IL-10+ (ANOVA=0.7101) subpopulations between treatment groups (**Fig. 5B&C**). In the spleen, there were no significant differences in the frequencies of CD11c+ CD103+ (ANOVA=0.3143) or CD11c+ CD103+ CD86+ (ANOVA=0.895) cells between groups (**Fig. 5D**). These data show that local factors likely mediate maintenance of tolerogenic dendritic cell phenotypes during pregnancy despite acute viral infection.

**FIG 5.**
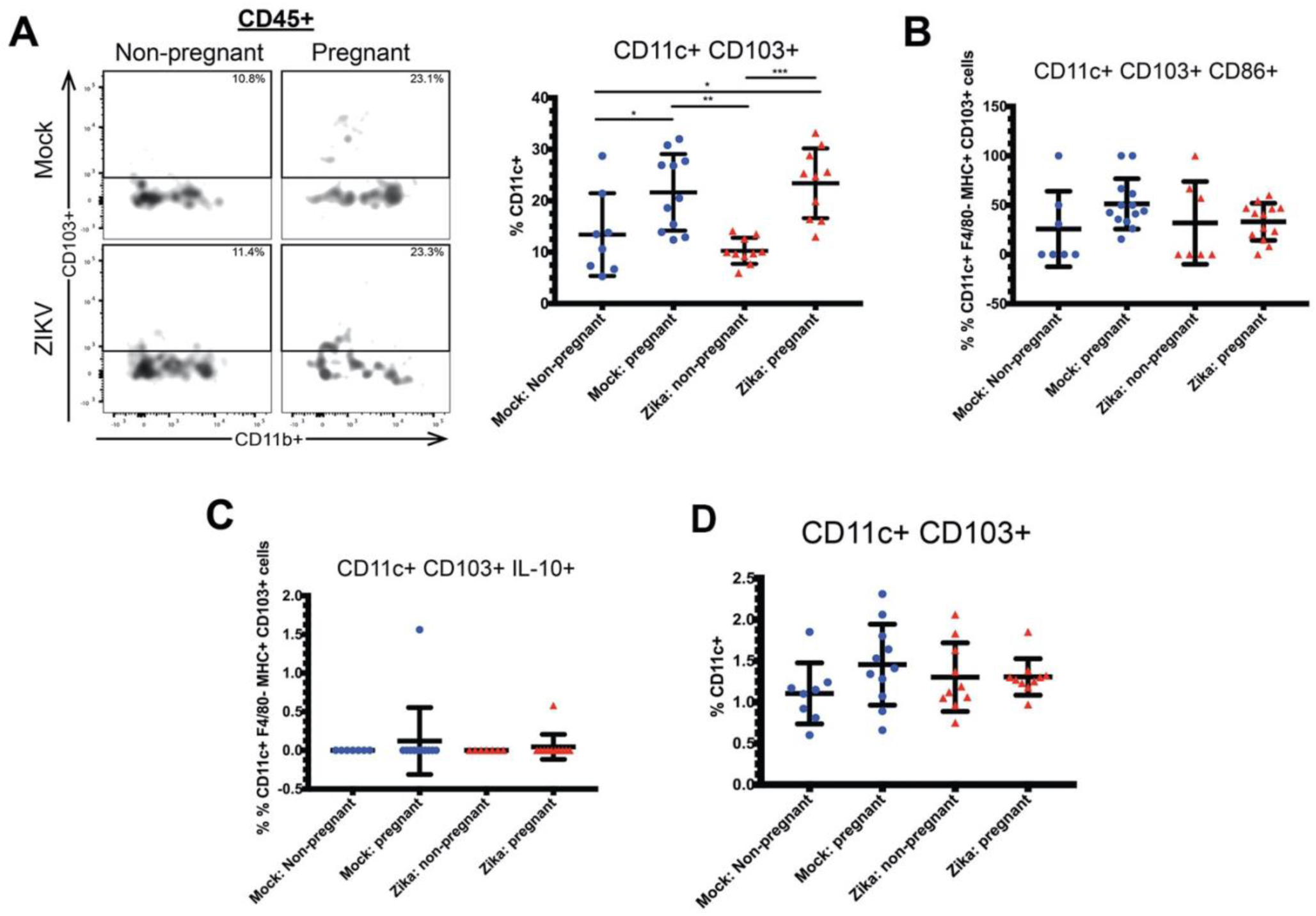
Pregnant mice have higher frequencies of CD11c+ CD103+ tolerogenic dendritic cells in the uterus. Flow cytometry was performed on the uterine immune cells of ZIKV-infected (red triangles) and mock-infected (blue circles) pregnant and non-pregnant mice. Frequencies of **A)** CD11c+ CD103+ cells, **B)** CD11c+ CD103+ CD86+ cells, and **C)** CD11c+ CD103+ IL-10+ cells were measured in the uterus. **D)** Frequencies of CD11c+CD103+ cells in the spleen. A representative flow cytometry plot (CD11c+ CD103+ cells within the CD45+ population) is shown in (A). *p<0.05, **p<0.01, ***p<0.001; one-way ANOVA and Tukey’s multiple comparisons test. N=40. Error bars represent the mean ± standard deviation.

### Pregnant mice exhibit decreased IL-12 responses to ZIKV infection in the uterus

Next, we evaluated the expression of pro-inflammatory cytokines in the uterine innate immune cells. IL-12 promotes the differentiation of T cells into Th1 cells and activates NK cells, and it is upregulated during certain viral infections (41, 42). Additionally, multiple studies have shown that IL-12 levels are increased in the blood and endometrial tissue of in women with recurrent pregnancy loss, suggesting that IL-12 may be detrimental during pregnancy (43, 44). In uterine tissue at day 3 post-infection, we found that the frequency of CD45+ cells producing IL-12 was decreased following ZIKV infection in pregnant compared to non-pregnant mice (**Fig. 6A**, p=0.0081, ANOVA= 0.007). This difference was not seen when non-pregnant and pregnant mock-infected mice were compared (p=0.9535). Next, we further analyzed IL-12 expression in several immune cell subtypes. We found that IL-12+ CD11b+ (**Fig. 6B,** p=0.007, ANOVA= 0.0059), IL-12+ CD68+ (**Fig. 6C**, p=0.0018, ANOVA= 0.0011), and IL-12+ Ly6C+ (**Fig. 6E**, p=0.004, ANOVA= 0.0028) cells were significantly decreased in ZIKV-infected, pregnant mice compared to ZIKV-infected non-pregnant mice. CD45+ CD103+ cells exhibited a trend towards decreased IL-12 expression during pregnancy in both mock and ZIKV-infected mice when compared to non-pregnant mice (ANOVA= 0.006) (**Fig. 6D**). There was no significant difference in CD11c+ IL-12+ cells between groups (data not shown, ANOVA=0.1943).

**FIG 6.**
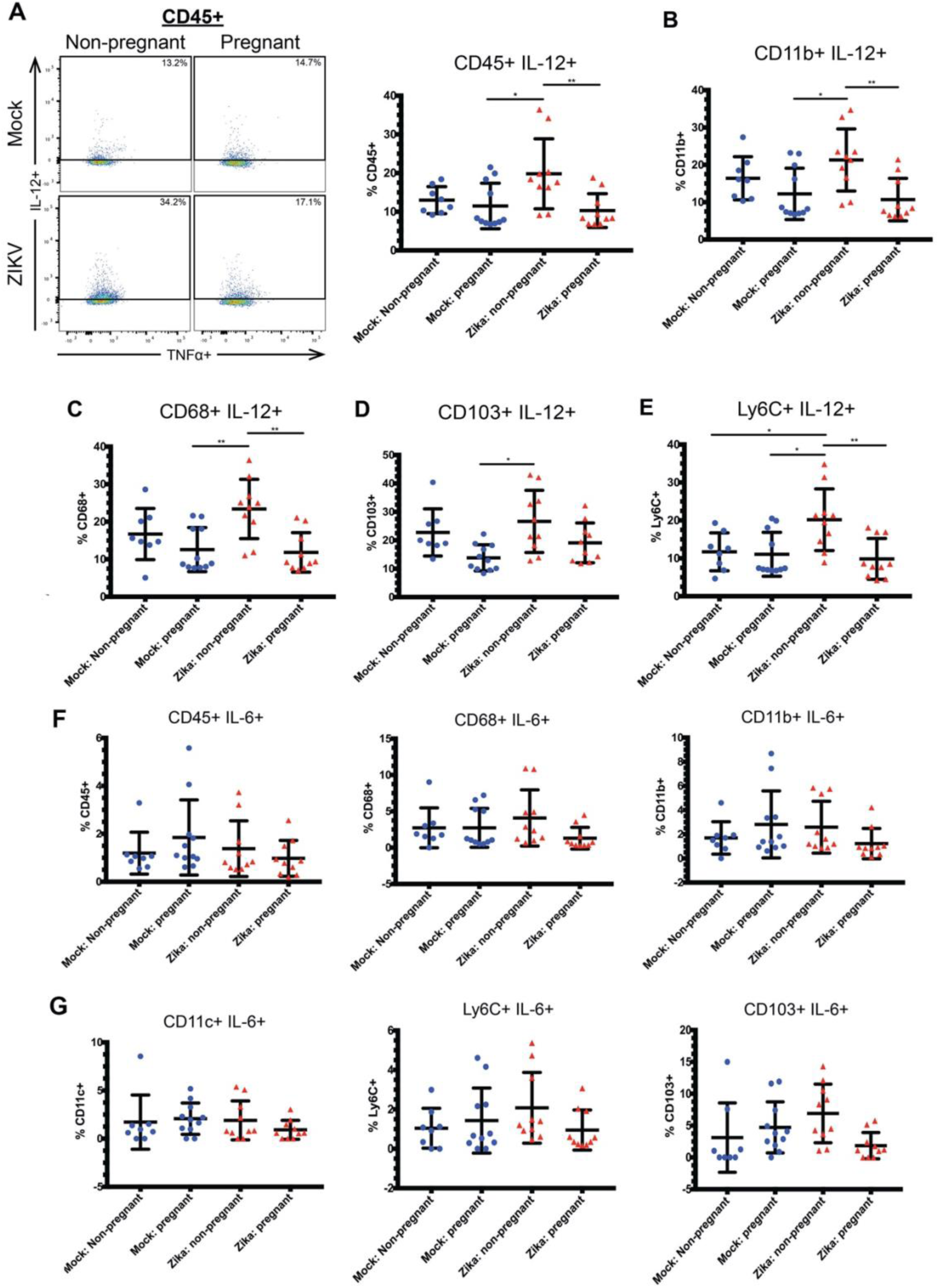
Pregnant mice have lessened IL-12 responses in the uterus during Zika virus infection. Flow cytometry was performed on the uterine immune cells of ZIKV-infected (red triangles) and mock-infected (blue circles) pregnant and non-pregnant mice. Frequencies of IL-12-expressing cells were measured within the uterine CD45+ (**A**), CD11b+ (**B**), CD68+ (**C**), CD103+ (**D**), and Ly6C+ (**E**) populations. (**F&G**) Frequencies of IL-6-expressing cells were measured within the uterine CD45+, CD68+, CD11b+, CD11c+, Ly6C+, and CD103+ populations. N=39. Representative flow cytometry plots depicting IL-12+ cells within the CD45+ population are shown in (A). *p<0.05, **p<0.01; one-way ANOVA and Tukey’s multiple comparisons test. N=39. Error bars represent the mean ± standard deviation.

Additionally, we measured the frequencies of IL-6+ cells within each of these populations, as it has been reported that IL-6 is upregulated during ZIKV infection (45). We found no significant differences in the frequencies of IL-6+ CD45+, IL-6+ CD68+, IL-6+ CD11b+, IL-6+ CD11c+, IL-6+ Ly6C+, or IL-6+ CD103+ cells between groups (**Fig. 6F&G**). These data show that pregnancy results in decreases in IL-12-expressing cells during ZIKV infection, chiefly IL-12+ monocytes and macrophages.

### Pregnant mice exhibit decreased IL-12 responses to ZIKV infection in the spleen

Our murine model utilized a localized intravaginal infection, and we found evidence of pregnancy-associated modulation of IL-12 expression in subsets of CD45+ cells in the uterus following infection. Next, we evaluated the spleen for similar changes in IL-12 expression. We found a significant decrease in splenic CD45+ IL-12+ cells in pregnant ZIKV-infected mice compared to non-pregnant ZIKV-infected mice (**Fig. 7A**, p=0.0323, ANOVA=0.0288). In the subset analysis, we found that frequencies of IL-12+ cells were significantly decreased within the CD68+ (**Fig. 7B**, p=0.0441, ANOVA=0.0241) and CD11b+ (**Fig. 7C**, p=0.0093, ANOVA=0.0107) populations in ZIKV-infected pregnant mice compared to ZIKV-infected non-pregnant mice. Similar to the uterus, inhibition of IL-12 expression appeared to be specific to monocytes and macrophages, as CD11c+ (**Fig. 7D**, ANOVA=0.2629) and CD103+ (**Fig. 7F**, ANOVA=0.9808) cells did not exhibit significant changes in IL-12 expression between treatment groups. The frequency of Ly6C+ IL-12+ cells trended toward a decrease in the ZIKV-infected pregnant mice compared to the other groups, but this change did not reach significance when the groups were compared to each other (**Fig. 7E**, ANOVA=0.0478). Similar to the results seen in the uterus, IL-6 expression did not differ significantly in any of the cell subsets tested (**Fig. 7G&H**).

**FIG 7.**
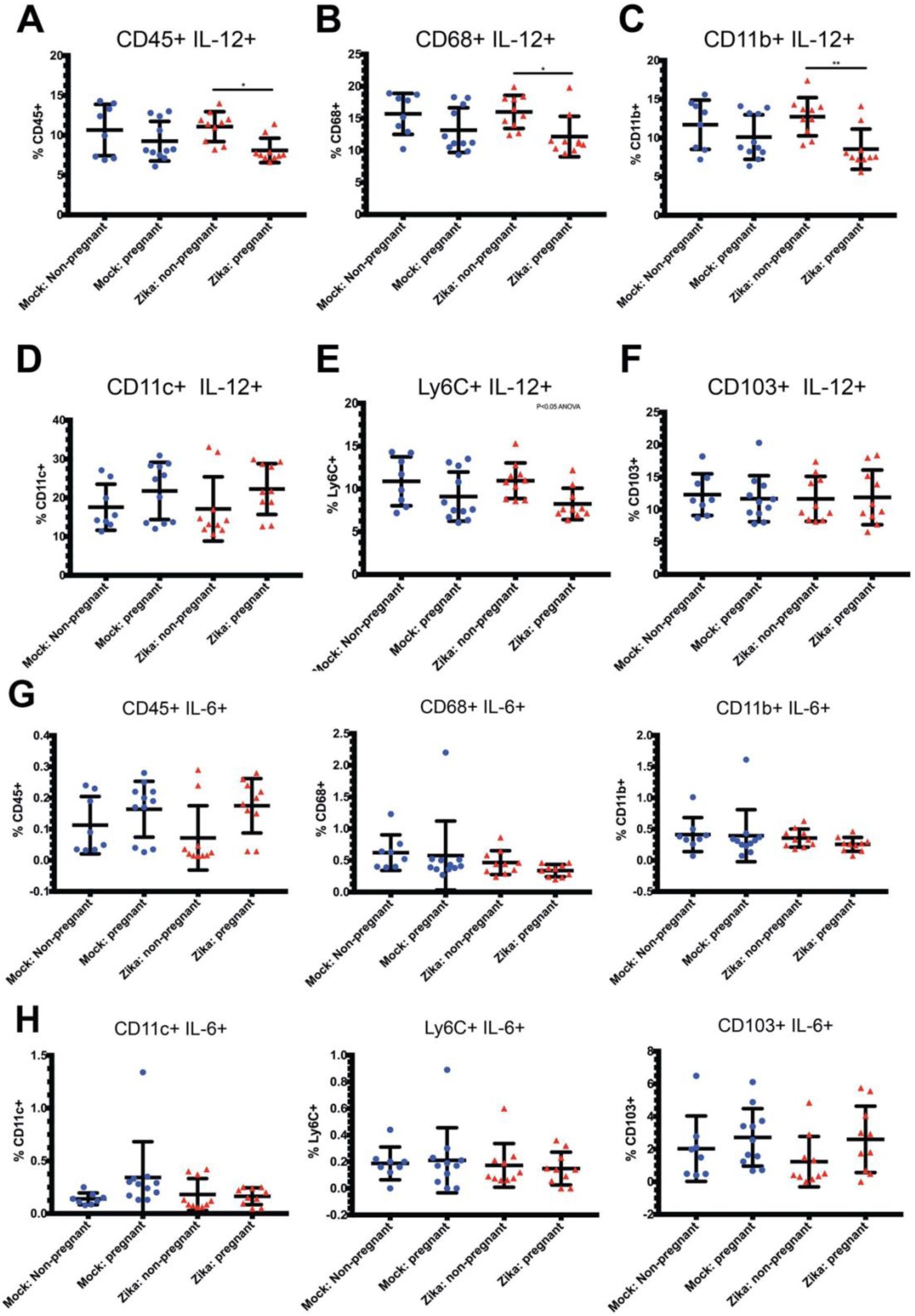
Pregnant mice exhibit decreased IL-12 responses in the spleen during Zika virus infection. Flow cytometry was performed on the splenic immune cells of ZIKV-infected (red triangles) and mock-infected (blue circles) pregnant and non-pregnant mice. Frequencies of IL-12-expressing cells were measured within the splenic CD45+ (**A**), CD68+ (**B**), CD11b+ (**C**), CD11c+ (**D**), Ly6C+ (**E**), and CD103+ (**F**) populations. (**G&H**) Frequencies of IL-6-expressing cells were measured within the uterine CD45+, CD68+, CD11b+, CD11c+, Ly6C+, and CD103+ populations. N=39. *p<0.05, **p<0.01; one-way ANOVA and Tukey’s multiple comparisons test. N=39. Error bars represent the mean ± standard deviation.

## Discussion

Our data are the first to evaluate acute, innate immune cellular responses in the genital tract of immune competent mice to Zika virus infection during early pregnancy. The data show that early stages of pregnancy results in inhibition of innate cellular activation and maintenance of tolerogenic immune cell changes despite acute viral infection in the genital tract. We show that pregnant mice exhibited decreased CD45+Ly6C+ cells in the spleen following acute viral infection. We also found that pregnancy induces expression of tolerogenic DCs (CD45+ CD11b+CD11c+) that express increased MHCII with ZIKV infection during pregnancy while decreasing expression of CD86. Moreover, IL-10 expression is significantly decreased during this early stage of pregnancy in mock infected animals likely representing the importance of some inflammatory responses to develop early pregnancy. However, during acute viral infection, IL-10 is not significantly suppressed in pregnant mice during ZIKV infection. This implies that ZIKV infection induces increased expression of tolerogenic signals (decreased CD86 and less suppression of IL-10) on CD11b+CD11c cells during pregnancy. These findings indicate that pregnancy inhibits virus-induced acute inflammation during early implantation likely as a mechanism to protect the developing fetus.

We also found that some of the inhibitory cell phenotypes were specific to the uterine tissue. For example, during acute ZIKV infection, non-pregnant mice significantly suppress immunotolerant CD11c+CD103+ cells to support acute inflammation for the infection; however, pregnant mice exhibit a significant increase in CD11c+CD103+ cells in the uterus. These changes were not seen in the spleen implying that the expression of CD11c+CD103+ cells in the uterus is regionally regulated to support the developing pregnancy.

In contrast, some of the inhibitory cell phenotypes were found in both the uterus and the spleen. We found that CD45+IL-12 responses to acute viral infection were significantly decreased in both the uterus and spleen of pregnant mice compared to non-pregnant mice. In the uterus, decreased IL-12 production was largely due to CD11b+, CD68+, and Ly6C+ cells, and in the spleen decreased IL-12 production was largely due to CD11b+ cells. These data show for the first time, that both systemic and regional responses during pregnancy modulate the acute, innate immune cellular response to acute viral infection.

In our mouse model, implantation occurs at E4 (46), and we inoculated mice intravaginally with ZIKV at this early stage of pregnancy. At this timepoint, embryo implantation is driving changes in immune cell infiltrates, which support the development of the pregnancy. Approximately 70% of decidual leukocytes are natural killer (NK) cells, 20-25% are macrophages, 1.7% are DCs, and approximately 3-7% are T cells (47, 48). The presence and modulation of each individual cell type is important to support decidual development and promote tolerance of the haploidentical fetus. However, the impact of the intricate immune modulation during pregnancy is not well examined during acute viral infection in the genital tract.

Macrophages are involved in remodeling of the spiral arteries during early pregnancy, a process which is crucial in establishing blood flow to the placenta (49, 50). Macrophages are broadly classified into two subtypes: classically activated M1 macrophages, which are considered pro-inflammatory, and alternatively activated M2 macrophages, which have anti-inflammatory properties and are involved in tissue repair (51-53). We found that pregnant mice exhibited higher frequencies of uterine CD68+ macrophages with lower expression of IL-12 upon ZIKV infection. Previous studies have shown that several M2 markers, including CD206, CCL18, CD163, IL-10, and mannose receptor c type (MRC)-1 are expressed on decidual macrophages (54-56). Human placental macrophages, or Hofbauer cells, are targets of ZIKV and promote dissemination of the virus. Infected Hofbauer cells produce pro-inflammatory cytokines, including MCP-1, IL-6, IP-10, and type I interferons (57). It is unclear how pregnancy-induced immunotolerance would impact responses of Hofbauer cells, and future studies should examine the interaction between maternal and placental immune regulation during acute viral infection.

CD11c+ DCs are crucial for early placentation and regulate tissue remodeling and angiogenesis (58). While the inflammatory activity of decidual DCs is important to support early implantation, they are altered by the local environment, resulting in loss of migration of uterine DCs to the lymph nodes (59). However, the specific changes in activation and cytokine production in DCs during acute infection have not yet been characterized. Tolerogenic DCs, including the CD11b+ CD11c+ and CD11c+ CD103+ populations, promote immunotolerance toward the fetus during pregnancy and secrete anti-inflammatory mediators including IL-10 (36-39). We found that both subtypes were upregulated in pregnant mice, regardless of infection. Interestingly, we found that there were fewer CD11b+ CD11c+ IL-10+ cells in the uteruses of mock-inoculated pregnant mice than mock-inoculated non-pregnant mice. However, in the ZIKV-infected pregnant mice did not decrease IL-10 expression or CD11c+ CD103+ cells. These results suggest that pregnancy signals maintain immunotolerant signaling and cell types despite acute viral infection in these tissues. Additionally, pregnant mice exhibited fewer CD86+ CD11b+ CD11c+ cells in the uterus, which is indicative of less mature, unactivated DCs. Taken together, these results indicate that pregnancy-associated tolerogenic DCs are not significantly suppressed by ZIKV infection. This likely impacts the induction of downstream immune responses to the virus. Further studies are needed to determine the effects of the pregnancy-induced tolerogenic immune environment on the anti-viral adaptive immune responses.

Our data also show that uterine monocytes, macrophages, and DCs are deficient in IL-12 production upon challenge with ZIKV during pregnancy. Since IL-12 is an important activator of NK cell responses(60), suppression of IL-12-induced activation of uterine NK cells is likely important to prevent NK activation and increased risk to the pregnancy. Our data show that virus-induced production of IL-12 by CD45+ cells is significantly reduced in pregnant mice compared to non-pregnant mice. This may be an important mechanism by which the localized immune response in the decidua is modulated to protect the pregnancy while still mounting an immune response to viral infection that is efficacious but not deleterious to the developing fetus. These findings should be evaluated as a potential biomarker of pregnancy loss during acute infection, as these markers may provide prognostic value for pregnancy loss and complications.

The findings of this study may be broadly applicable to other acute infections during pregnancy, and further studies are needed to evaluate pregnancy-induced immune modulation during acute infection and vaccination. Additionally, our data show that activation of important antigen presenting cells are modulated during pregnancy.. These findings have important implications for vaccine studies during pregnancy as well, since dendritic cells and other antigen presenting cells are vital for development of the adaptive immune response that defines vaccine outcomes. In conclusion, our results show that pregnancy-induced immunotolerance impacts the acute innate cellular response to ZIKV infection and inhibits important features of the acute anti-viral immune response. Further studies are also needed to examine the impact of pregnancy-induced modulation of acute anti-viral immune responses on the adaptive immune response, in pregnancy outcomes during acute infection, and in vaccine outcomes during pregnancy.

## Materials and Methods

### Ethics Statement

All animal research was approved by the University of Colorado and Denver VAMC local Institutional Animal Care and Use Committees. Approval number 1098v3. All laws and regulations regarding animal care and euthanasia were followed according to guidelines from the PHS/NIH/OLAW policy, Animal Care Policy (USDA), and the AVMA guidelines on euthanasia.

### Virus propagation and cell culture

Vero cells (ATCC, Manassas, VA) and C6/36 cells (ATCC) were cultured at 37°C and 5% CO_2_ in complete Minimal Essential Media (MEM) supplemented with 10% fetal bovine serum (FBS, HyClone, Thermo Fisher Scientific, Waltham, MA). ZIKV strain PRVABC59 (GenBank: KU501215) was provided by the Centers for Disease Control (CDC, Atlanta, GA). ZIKV stocks were propagated in Vero cells at passage 4 and C6/36 cells at passage 1, and cell culture supernatants were harvested at 6 days post infection. Virus stocks were titrated in Vero cells using a focus forming assay (FFA) and were aliquoted and stored at −80°C.

### Mice

Six-week-old C57BL/6J (stock no. 000664) male and female mice were purchased from Jackson Laboratory (Bar Harbor, ME). The mice were housed in a pathogen-free animal facility at the University of Colorado Anschutz Medical Campus (Aurora, CO) and maintained on a 12:12 light/dark cycle at 21-24°C. Eight-week-old female mice were mated with male mice ranging from 8-20-weeks-old. Each mating pair was housed separately.

### Hormone Treatment

To increase the likelihood of pregnancy in the mice, female mice were treated with exogenous gonadotropins to increase ovulation (61). Two days before mating, female mice were injected intraperitoneally with 2.5 iu of Pregnant Mare Serum Gonadotropin (bioWORLD, Dublin, OH). 48 hours later, they were intraperitoneally injected with 2.5 iu human chorionic gonadotropin (HCG, Sigma Aldrich, St. Louis, MO) and immediately mated with male mice overnight (16 hours). The males and females were separated the following morning (E0.5).

### Zika virus infection

On day E4.5, the eight-week-old pregnant and non-pregnant female mice were randomly assigned to either the mock or ZIKV infection groups. The mice were anaesthetized with isoflurane (McKesson Corporation, Irving, TX) and infected intravaginally with 5×10^5^ FFU of PRVABC59 ZIKV in 15 µL of HBSS (Gibco, Thermo Fisher). Mock-infected mice received 15 µL of HBSS intravaginally. Spleen and uterine tissues were harvested for flow cytometry at 3 days post infection (E7.5).

### Vaginal lavages

Vaginal lavages were performed 48 hours post infection. Mice were anaesthetized with isoflurane, and 50 µL of sterile phosphate buffered saline (PBS, Corning, Corning, NY) was inserted into the urogenital tract using a micropipette. The liquid was expelled slowly into the urogenital tract and then drawn back up and mixed with 200 µL of sterile PBS supplemented with 1% FBS. The samples were vortexed for 30 seconds and then aliquoted and stored at −80°C.

### RNA extraction and qPCR

ZIKV RNA was isolated from the vaginal lavage samples using the E.Z.N.A. Viral RNA Kit (Omega Bio-tek, Norcross, GA) according to the manufacturer’s instructions. The primer and probe set Zika1087/1108FAM/1163c (IDT, Coralville, Iowa) was used to detect viral RNA. Real-time qPCR was performed using the Luna Universal Probe qPCR Master Mix (New England Biolabs, Ipswich, MA) with amplification on the Biorad CFX96 Real Time PCR Detection system, both per the manufacturer’s instructions. The sensitivity of this assay was evaluated by testing known dilutions of an RNA transcript copy of the ZIKV P1 plasmid. Concentration of viral RNA (copies/microliter) was calculated using the standard curve generated by the CFX96 instrument.

### Tissue processing

Spleen tissues were processed into single-cell suspensions by mechanical dissociation; tissues were crushed through a 70 µm cell strainer (CELLTREAT, Pepperell, MA) using disposable plastic pestles (CELLTREAT). Red blood cells (RBC) were removed by incubating the cell suspensions in 5 mL 1X RBC Lysis Buffer (eBioscience, Thermo Fisher) for 5 minutes at room temperature. The cells were then washed in 30 mL of R10 media (RPMI with L-glutamine (Corning) + 10% FBS + 1% Penicillin/Streptomycin (Corning) + 1% HEPES (Gibco) + 1% Sodium Pyruvate (Gibco) + 1% MEM Non-essential amino acids (MEM-NEAA, Gibco)), vortexed, and centrifuged at 500 rcf.

Uterine tissues were enzymatically digested using Liberase TL (Roche, Basel, Switzerland) at a final concentration of 160 µg/mL in HBSS (Gibco, Thermo Fisher). First, each tissue was suspended in 500 µL of cold liberase + HBSS in a 1.5 mL Eppendorf tube and mechanically dissociated using small surgical scissors. Next, another 500 µL of Liberase + HBSS was added to each tissue, and the samples were incubated at 37°C for 35 minutes with occasional vortexing. Samples were kept on ice in between steps. After incubation at 37°C, the dissociated tissues were filtered through 100 µm cell strainers (CELLTREAT).

After preparation of single-cell suspensions, the samples were centrifuged at 500 rcf for 5 minutes, counted using Trypan blue (Corning), and resuspended at a concentration of 1×10^6^ cells/mL in R10 media. The cells were then aliquoted into FACS tubes (0.25-1×10^6^ cells/tube) with strainer caps (BD Biosciences, San Jose, CA).

### Flow cytometry

The following antibodies were used for extracellular flow cytometry: anti-mouse CD45 BV650 (clone 30-F11, Biolegend, San Diego, CA), anti-mouse/human CD11b APC-Cy7 (clone M1/70, Biolegend), anti-mouse CD11c PE-eFluor 610 (clone N418, eBioscience), anti-mouse I-A/I-E (MHCII) FITC (clone M5/114.15.2, Biolegend), anti-mouse CD103 BV711 (clone 2E7, Biolegend), anti-mouse Ly6C BV785 (clone HK1.4, Biolegend), anti-mouse CD24 PerCP-eFluor 710 (clone M1/69, eBioscience), anti-mouse CD86 PE-Cy7 (clone GL-1, Tonbo Biosciences, San Diego, CA), and anti-mouse F4/80 BUV395 (clone T45-2342, BD Biosciences). The following antibodies were used for intracellular flow cytometry: anti-mouse CD68 PE-Cy7 (clone FA-11, Biolegend), anti-mouse CD3 BUV395 (clone 145-2C11, BD Biosciences), anti-mouse CD3 BV785 (clone 17A2, Biolegend), anti-mouse IL-12 (p40/p70) PE (clone C15.6, BD Biosciences), anti-mouse IL-6 APC (clone MP5-20F3, BD Biosciences), and anti-mouse IL-10 APC (clone JES5-16E3, Biolegend). Ghost Violet 510 dye (Tonbo Biosciences) was used to assess viability.

Single cell suspensions were washed in PBS, centrifuged at 500 rcf for 5 minutes, and briefly vortexed. Next, 10 µL of viability dye (0.1 µL dye + 10 uL FACS buffer (1% FBS in PBS) per sample) was added to each sample, vortexed, and incubated at room temperature for 10 minutes. Next, 50 µL of extracellular antibodies prepared in FACS buffer were added to each sample, vortexed, and incubated at 4°C for 25 minutes. 210 µL of Cytofix/Cytoperm solution (BD Biosciences) per sample was then added to permeabilize the cells, followed by vortexing and incubation for 20 minutes at 4°C. The cells were then washed in 1 mL of 1x Perm/Wash buffer (BD) or Flow Cytometry Perm Buffer (Tonbo Biosciences) twice, centrifuged at 700 rcf for 5 minutes, and vortexed. Next, 50 µL of intracellular antibodies in Perm/Wash or Perm buffer were added to each sample, vortexed, and incubated at 4°C of 45 minutes. The samples were then washed once more in Perm/Wash or Perm buffer, centrifuged at 700 rcf for 5 minutes, vortexed, and finally fixed in 1% paraformaldehyde (Thermo Fisher).

The data was acquired on a LSRII flow cytometer (BD) using voltages standardized according to previously published methods (62). FlowJo software (FlowJo, LLC, Ashland, Oregon) was used to analyze the data. The gating strategies

### Statistics

All statistical analysis was performed in Prism 7 software (GraphPad, San Diego, CA). One-way ANOVA and Tukey’s multiple comparison tests were used to compare cell frequencies between pregnant and non-pregnant mock and ZIKV-infected mice. P values, F values, and degrees of freedom for each parameter measured are shown. T tests were used when only two groups were compared. All the data are presented as mean ± standard deviation. P<0.05 was considered statistically significant. All data shown represent 2 experiments of 19-20 mice each (n=39-40 mice total for each parameter measured).

## Acknowledgements

Funding for this study was provided by the United States Department of Defense grant (PRMRP PR160117), VA Merit award I01BX003863, and CCTSI Co-Pilot CU/CSU Award to J.D.B.

The authors would like to thank Hadrian Sparks, Monica Graham, Brendan Monogue, and Aaron Massey for important discussions regarding development of the data and manuscript.

